# Loss of short-form Ron promotes B and T cell cooperation to prevent outgrowth of breast cancer bone metastasis

**DOI:** 10.64898/2026.09.05.749628

**Authors:** Clint H. Valencia, Marija Nadjsombati, Truc Pham, Danny Soltero, Navina Devarajan, Jaime Fornetti, Alana L. Welm

## Abstract

While immunotherapy has great potential to eradicate tumors, success in metastatic breast cancer is limited. Bone metastasis, the most common site for breast cancer metastasis, has been associated with weakened anti-tumor immunity, both in the bone and in other metastatic sites. Additional strategies promoting anti-tumor immunity are needed. Using the MMTV-PyMT intratibial model of breast cancer bone metastasis, we discovered that deletion of host short-form Ron (Ron SF^-/-^) protected against bone metastatic outgrowth by promoting strong infiltration of lymphocytes, including B cells, into tumors at the micrometastasis stage. Transcriptionally, Ron SF^-/-^ B cells were more active and inflammatory compared to wildtype. Elimination of mature B cells (JH^-/-^) rescued tumor growth in Ron SF^-/-^ mice, indicating that mature B cells were necessary for anti-tumor immunity and protection against bone metastasis. In Ron SF^-/-^ hosts, T follicular helper (Tfh) cells were abundant in tumor-enriched regions of bone, suggesting interactions between B and T cells. B-T cell cooperation was further supported by attenuation of CD4 and CD8 T cell tumor infiltration in Ron SF^-/-^JH^-/-^ hosts and, conversely, diminished B cell tumor infiltration and rescue of tumor growth in Ron SF^-/-^ hosts lacking CD4 or CD8 T cells. B cells were not required to protect Ron SF^-/-^ mice from lung metastasis, indicating a tissue-dependent anti-tumor role for B cells. These data support a positive feedback loop between B and T cells regulated by SF-Ron and reveal the anti-tumor potential of B cells in the bone.

## Introduction

In the United States, there are more than 300,000 new diagnoses and 40,000 deaths annually from breast cancer (1). Despite improved treatments and longer survival times, many still die when cancer cells metastasize to organs such as the bones, lungs, brain, or liver (2). Most metastases are detected years after the primary tumor is resected, when the cancer recurs (2,3). In many cases, the long interval between initial diagnosis and recurrence is attributed to disseminated tumor cells remaining dormant in tissues such as bone or persisting as micrometastases before metastatic outgrowth (3).

The bone is the most common site of breast cancer metastasis; about 70 percent of metastatic breast cancer patients have bone metastasis (4–7). Bone metastasis profoundly affects patients by increasing their risk of skeletal fractures and is frequently followed by metastatic progression to extraosseous sites (6,8,9). Breast cancer bone metastases are predominantly osteolytic, where bone destruction results from disruption of bone homeostasis normally maintained by a balance of osteoclast and osteoblast function (4). Current treatments for bone metastasis aim to protect bones by antagonizing osteoclast activity with bisphosphonates or the RANKL antagonist denosumab, typically in combination with systemic therapies and local treatments such as surgery and/or radiation (6,7,10). Despite current therapies, patients with bone metastases continue to experience severe skeletal complications and subsequent metastatic progression to other sites (8,9,11), highlighting the need for new strategies to treat or prevent bone metastasis.

One strategy to reduce deaths from breast cancer is to improve the immune response against tumors to effectively prevent metastatic disease. For successful immune-mediated tumor control, infiltration of immune cells into tumors is vital, and patients with high immune cell infiltration into their tumors have better responses to immunotherapy and improved outcomes (12,13). In breast cancer, most metastatic tumors have low immune cell infiltration and are poorly responsive to immunotherapies (12,13). Bone metastases, in particular, have an immunosuppressive microenvironment and can attenuate the systemic response to immunotherapy in multiple cancer types, including breast (14–16). This immunosuppressive nature and the systemic consequences of breast cancer bone metastasis, as well as the high frequency of bone metastasis in patients, make it imperative to find new ways to control the disease in this site.

We and others have reported the receptor tyrosine kinase Ron as a key regulator of cancer metastasis, where it functions in both tumor cells and in the immune system (17–20). The gene encoding Ron gives rise to two transcripts: full-length Ron (FL-Ron) and short-form Ron (SF-Ron) (18). FL-Ron and SF-Ron share the same kinase domain, and the gene is overexpressed in several cancers, including breast cancer (17,21). SF-Ron is transcribed from an alternative promoter within intron 10 of the *Stk* gene in mice or within exon 10 of the *MST1R* gene in humans and is reported to encode a truncated, constitutively active isoform of Ron kinase (18,22). When overexpressed in tumors, either isoform enhances tumor progression and metastasis (21,23–25). In the host, FL-Ron activation in tissue-resident macrophages by its ligand, macrophage-stimulating protein, skews them toward a tumor-promoting anti-inflammatory (M2) phenotype (26,27). Additionally, mice that lack all Ron kinase activity fail to downregulate inflammatory responses when challenged with infection or injury to the liver or lungs (28,29), further supporting a role for Ron in immune regulation.

While FL-Ron has been extensively studied in tumors and host cells (23), SF-Ron has been severely understudied. Although both isoforms can function similarly, it has been suggested that FL-Ron and SF-Ron play divergent roles in tumors and noncancer-related inflammation (21,23,30). In the tumor setting, our lab has shown that mice lacking all Ron tyrosine kinase activity (19,20) are protected from breast cancer lung metastasis due to stronger anti-tumor immune responses. This is recapitulated in mice lacking only SF-Ron (31), indicating that loss of host SF-Ron alone is sufficient to promote anti-tumor immunity in these models. These findings support a non-redundant role for SF-Ron that is distinct from FL-Ron.

We have also reported a role for Ron in bone metastasis models, in which loss of host Ron tyrosine kinase activity protected mice from osteoporosis and tumor-induced osteolysis by reducing osteoclast resorption activity (32). However, the specific role of SF-Ron in bone metastasis or bone tumor immunity remains unknown. Thus, we sought to determine whether SF-Ron regulates the anti-tumor immune response in other metastatic sites, such as breast cancer bone metastases. Here, we report that loss of SF-Ron promotes a strong tissue-specific anti-tumor immune response through B cell and T cell cooperation within bone micrometastases, resulting in tumor clearance and protection against overt bone metastasis.

## Materials and Methods

### Animals

All procedures involving mice were approved by the University of Utah Institutional Animal Care and Use Committee (IACUC). Our studies focused on 6- to 8-week-old female animals to prioritize investigating female breast cancer. Ron SF^-/-^ mice were previously described (30). All mice for this study were backcrossed to the FVB genetic background and confirmed to be 98-99% FVB using the MiniMUGA genotyping array from Transnetyx. Wild-type (WT) FVB/NJ mice were purchased from The Jackson Laboratory (Strain #001800) and bred in-house. To generate F1 mice for the FVB strains CD8^-/-^ (33), CD4^-/-^ (33), or JH^-/-^ (34), cryopreserved sperm from fully homozygous mice were a generous gift from Dr. Lisa Coussens and used for in vitro fertilization of WT FVB/NJ oocytes. F1 offspring were heterozygous for the allele of interest and intercrossed to generate F2 progeny with the expected Mendelian distribution. Homozygous breeding pairs were prioritized in subsequent generations (F3–F5) to establish a fully homozygous colony. Tail DNA for routine genotyping was isolated in 100 mmol/L tris(hydroxymethyl)aminomethane (Tris), 5 mmol/L ethylenediaminetetraacetic acid (EDTA), 0.2% sodium dodecyl sulfate (SDS), 200 mmol/L sodium chloride (NaCl) in deionized water with 100 µg/mL proteinase K, and PCR was performed (primers are listed in Supplementary Table S1). Each CD8^-/-^, CD4^-/-^, and JH^-/-^ strain was then crossed with Ron SF^-/-^ mice to generate homozygous double knockouts.

### Tumor injections and tissue processing

Pooled mammary tumor cell stocks were obtained from Mouse Mammary Tumor Virus - Polyoma Virus middle T antigen (MMTV-PyMT) mice on the FVB/NJ background (35). Two days prior to injection, tumor cells were thawed and cultured in Dulbecco’s Modified Eagle Medium/Nutrient Mixture F-12 (DMEM/F12) medium (Gibco, Invitrogen) supplemented with fetal bovine serum (FBS) (10%; Gibco, Invitrogen), insulin–transferrin–selenium–ethanolamine (1×; Gibco, Invitrogen), recombinant murine epidermal growth factor (EGF) (10ng/mL; Invitrogen), hydrocortisone (1 μg/mL; Sigma), penicillin–streptomycin (1x; Gibco, Invitrogen), and gentamicin (50 μg/mL; Apex, Genesee Scientific). For intratibial (IT) injections, 1 × 10^5^ MMTV-PyMT tumor cells in 10µL Matrigel (Corning) were injected into the right tibia of anesthetized female mice. At the endpoint, mice were euthanized and hindlimbs collected. For histology, bones were fixed in 10% neutral-buffered formalin for 24-48 hours at room temperature, decalcified in Formical-2000 (StatLab) for 7-8 days at 4°C, and stored refrigerated in 70% EtOH until paraffin embedding. Tumors were confirmed by immunohistochemistry (IHC) for EpCAM (below). Experimental lung metastasis via tail-vein (TV) injection was performed as previously described (31). For flow cytometry, bulk RNA-sequencing (RNA-seq), and single-cell RNA-sequencing (scRNA-seq) on cells from the bone, bones were gently crushed in media (1x phosphate-buffered saline (PBS), 2% FBS, 0.5 mM ethylenediaminetetraacetic acid (EDTA)) using a mortar and pestle, and bone marrow was isolated by pipetting. For tumor-enriched areas, only the region of the bone with a visibly white marrow cavity (indicating that the normal marrow had been replaced by tumor cells) was used. For marrow, contaminating red blood cells were lysed using ammonium–chloride–potassium (ACK) lysis buffer and filtered through 100-μm nylon mesh to obtain single-cell suspensions.

### Ex vivo radiography and analysis

Prior to decalcification, osteolysis was assessed on fixed tibiae ex vivo using a Hologic/Faxitron UltraFocus DXA (51 kV for 6.3 sec). Osteolytic lesions were identified as foci of increased radio translucency within the injected tibia. The osteolytic lesion area was quantified manually using ImageJ-based FIJI software (36) (Supplementary Fig. S1B). Results are reported as a percentage lytic area (osteolytic area/whole tibia area).

### Immunohistochemistry (IHC) and quantification

Four-micron-thick paraffin sections were baked at 60°C for 1 hour, deparaffinized in CitriSolv (Decon Labs), and rehydrated in serial ethanol dilutions. Heat-induced epitope retrieval (HIER) in 10 mmol/L sodium citrate (pH 6.0) was performed overnight in a 60°C water bath. Endogenous peroxidase activity was quenched with 3% hydrogen peroxide in methanol for 10 minutes, followed by nonspecific blocking for one hour in PBS containing 5% bovine serum albumin (BSA), 10% normal goat serum, and 10% normal human serum, and mouse FcReceptor blocking reagent (Miltenyi Biotec; 1:100). After blocking, the sections were incubated with the primary antibody overnight at 4°C or for 1 hour at room temperature, followed by EnVision+ goat anti-rabbit horseradish peroxidase (HRP) polymer reagent (Agilent Technologies) or ImmPRESS HRP goat anti-rat IgG (Vector Laboratories), according to the species of primary antibody. Chromogen detection was performed using the 3,3′-diaminobenzidine kit (Vector Laboratories), and tissues were counterstained with Mayer’s hematoxylin (Sigma-Aldrich).

Following dehydration and clearing, sections were mounted using Cytoseal 60 (Epredia). Primary mouse antibodies used for histology analysis in this study were all unconjugated and include CD3e (clone D4V8L; Cell Signaling Technology; 1:100), CD4 (clone D7D2Z; Cell Signaling; 1:50), CD8a (clone D4W2Z; Cell Signaling; 1:250), B220 (CD45R; clone RA2–6B2; Santa Cruz Biotechnology; 1:800), EpCAM (clone D9S3P; Cell Signaling; 1:200), and CD11c (clone D1V9Y; Cell Signaling; 1:100). EpCAM primary antibody was incubated for 1 hour at room temperature, while the rest of the primary antibodies were incubated overnight at 4°C.

Image acquisition was performed at 20× magnification using whole-slide scanners: the 3D Histech Panoramic MIDI or the Zeiss Axioscan Z1. Tumor burden was quantified by dividing the EpCAM+ tumor cell area by the total marrow area. Results are represented as a percentage (tumor area/marrow area). At the 4-week timepoint, experiments showed that most Ron SF^-/-^ bones had no tumors, so results are reported as the fraction of mice with tumors rather than the percentage of marrow occupied by tumors. In the lung metastasis model, Ron SF^-/-^ hosts have very few tumors 4 weeks post-TV injection, so results are reported as the fraction of mice protected from tumor outgrowth. Quantification of the percentage of lung area occupied by metastasis was performed as previously described (31). Immune cell quantification was performed on five to ten biological replicates per group and restricted to tumor-bearing regions of the bone marrow cavity. Computer-assisted quantification of EpCAM, CD3e, B220, CD11c, CD4, or CD8a from images was performed in QuPath (37) using a standard threshold for each marker across all images.

### Flow cytometric analysis

Single-cell suspensions, described above, were stained with fixable viability dye in PBS for 30 minutes on ice, then incubated with anti-mouse CD16/32 Fc receptor-blocking antibodies (Clone 93; BioLegend; 1:200) for 10 minutes on ice. For surface marker staining, cells were stained with fluorophore-conjugated antibodies in Brilliant Stain Buffer (BD Biosciences) for 30 minutes on ice. The stained samples were acquired on a BD LSRFortessa flow cytometer, and immune population analyses were performed in FlowJo. Approximately 3 x 10^5^ events were collected for each sample. Forward-scatter versus side-scatter gating was first used to omit cellular debris and doublets. The single cells were then gated to eliminate dead cells, followed by gating on specific immune subsets (Supplementary Fig. S2A). The following mouse antibodies and stains were used: CD45-BUV496 (clone 30-F11; BD Biosciences; 1:200), CD3e-BUV395 (clone 145-2C11; BD Biosciences; 1:200), B220-PE-CF594 (clone RA3-6B2; BD Biosciences; 1:400), CD19-APC (clone 6D5; BioLegend; 1:200) or -Alexa Fluor 700 (clone 6D5; BioLegend; 1:200), CD11c-BV711 (clone N418; BioLegend; 1:300), CD335 (NKp46)-BV650 (clone 29A1.4; BioLegend; 1:200), CD11b-BV510 (clone M1/70; BD Biosciences; 1:300), F4/80-PE-Cy5 (clone BM8; eBioscience; 1:300), Ly6G-PE (clone 1A8-Ly6g; eBioscience; 1:300), Ly6C-PerCP-Cy5.5 (clone HK1.4; eBioscience; 1:300), CD4-BV86 (clone GK1.5; BD Biosciences; 1:400), CD8a-Alexa Fluor 700 (clone 53-6.7; BioLegend; 1:300) or -PE-Cy7 (clone 53-6.7; Cytek; 1:400), FOXP3-PE (clone FJK-16s; eBioscience; 1:100), CD185 (CXCR5)-BV421 (clone L138D7; BioLegend; 1:200), CD279 (PD-1)-APC (clone RMP1-30; eBioscience; 1:200), Ghost Dye Red 780 (Cytek; 1:4000).

### NK cell and plasmacytoid dendritic cell depletion

For NK cell depletion, mice were injected with 100 μg anti-mouse NK1.1 (clone PK136; BioXCell) or immunoglobulin G2a isotype control antibody (clone C1.18.4; BioXCell) in 200 μL of InVivoPure pH 7.0 Dilution Buffer (BioXCell). A single dose of antibody was injected intraperitoneally 2 days before tumor cell injection and once weekly post-injection (38). For plasmacytoid dendritic cell (pDC) depletion, 250 μg anti-mouse CD317 (BST2) (clone 927; BioXCell) or immunoglobulin G2b isotype control antibody (clone LTF2; BioXCell) in 200 μL of InVivoPure pH 7.0 Dilution Buffer (BioXCell) was injected intraperitoneally once daily for 2 days, and tumor cells were injected on day 3 (39). Post-tumor cell injection, mice were dosed every 2 days until endpoint. For both studies, mice were euthanized 4 weeks post-intratibial injection, and metastatic tumor burden was quantified as described above.

### Bulk RNA sequencing and analysis

B220+ cells were purified by using fluorescence-activated cell sorting (FACS) from tumor-enriched bone marrow regions of WT (n=3) and Ron SF^-/-^ mice (n=3) using fluorescence-activated cell sorting (FACS). Each single n consists of 3 pooled mice from each genotype to increase the number of B220+ cells collected. RNA was isolated from 5 x 10^5^ to 1 x 10^6^ B220+ cells using the Qiagen RNeasy Plus Mini Kit. Libraries were prepared using NEB Next Ultra II Directional Library Prep with rRNA depletion kit and sequenced on the Illumina NovaSeqX, to a depth of 33 million reads. Read counts were normalized using the rlog method in the DESeq2 R package, version 1.50.2, filtering genes with ≤5 reads. Differential expression analyses were also performed using DESeq2. Subsequent analyses were conducted using the R Studio environment, version 2026.7.1.147. Heatmaps were generated using the R package “hciR” version 1.8. Principal component analysis (PCA) plots and volcano plots were generated using the R package “ggplot2” version 4.0.3, with -log10 (Padj) > 1.3 considered significant. Gene set enrichment analysis (GSEA) was performed using GSEA software (version 4.4.0). The total gene list was preranked using log2 fold change (FC); the analysis was conducted with 1,000 permutations and “no collapse.” Defined gene signatures were selected from the Molecular Signatures Database, selecting mouse Hallmark and Gene Ontology as reference databases. For GSEA, P value < 0.05, FDR <0.25, and NES < -1 or >1 were considered significant. Bar plots were produced using GraphPad Prism version 11.

### Single-cell RNA sequencing (scRNA-seq) and analysis

FACS-purified B220+ cells from tumor-enriched regions of WT and Ron SF^-/-^ bones (5 WT and 5 Ron SF^-/-^; approximately 5,000 cells/sample) were processed with droplet-based 5′-end scRNA-seq using the GEM-X Universal 5′ Gene Expression V3 4-plex kit (10× Genomics) according to the manufacturer’s protocol, enabling multiplexing. Dual-indexed libraries were sequenced on the Illumina NovaSeqX with a depth of 30,000 reads per cell. Sequence reads were preprocessed using the 10× Genomics CellRanger pipeline to exclude mitochondrial genes and account for variance in unique molecular identifier counts, then further analyzed with the R package “Seurat” version 5.5.0 (40). The 10 data sets, 5 WT and 5 Ron SF^−/−^, were integrated into a combined data set. The Seurat pipeline was then applied to the combined dataset. Principal component analysis (PCA) and Uniform Manifold Approximation and Projection (UMAP) were performed using the first 10 PCA components. Cell clusters were identified using Seurat’s FindClusters function, and a resolution of 0.1 was used for all analyses. To examine B cell subsets, clusters expressing *Cd19*, *Cd79a*, *Cd79b, and Ms4a1* were extracted from the combined data set and re-clustered using 10 PCA components. The mean marker expression for each cluster was used to generate the dot plot. The biological identities of cell clusters were annotated by marker expression and using the web-based CIPR tool, which compares the cell cluster signatures with the publicly available Immunological Genome Project (ImmGen) database (41). UMAPs were produced using the R package “Seurat” version 5.5.0. Differentially expressed genes were identified using pseudobulk analysis in the R package “DESeq2” version 1.50.2, with raw counts aggregated per sample per cluster via Seurat’s AggregateExpression function. Dot plots were generated using Seurat’s DotPlot function. Heatmaps were created using the R package “pheatmap” version 1.0.13. Volcano plots were generated using the R package “ggplot2” version 4.0.3, with -log10 (Padj) > 1.3 considered significant. Gene set enrichment analysis (GSEA) was performed using GSEA software (version 4.4.0). The total gene list was preranked using log2 FC; the analysis was conducted with 1,000 permutations and “no collapse.” Defined gene signatures were selected from the Molecular Signatures Database, selecting the mouse Hallmark and Immunological Signatures as reference databases. For GSEA, P value < 0.05, FDR <0.25, and NES < -1 or >1 were considered significant. Bar plots were produced using GraphPad Prism version 11.

### Statistical analysis

For animal studies, sample size was calculated based on previous data to correctly detect differences of 20% or greater between groups (10% significance level and 80% power). Values are expressed as mean ± SEM of biological replicates. Presence of tumors between groups was compared using Fisher’s Exact Test. One-way ANOVA was used to compare lytic areas, followed by Tukey’s multiple comparisons test. An unpaired two-tailed multiple Student *t* test was used to compare unmatched groups with Gaussian distributions. Statistics were determined using GraphPad Prism version 11. P ≤ 0.05 was considered significant.

### Code availability

The analyses in this study were performed using publicly available R packages and established coding methods, as described in the methods section. No custom software or code was developed for this study. R scripts used for data analysis are available from the corresponding author upon request.

### Data availability

The data generated from bulk RNA sequencing and scRNA sequencing in this study are deposited in Gene Expression Omnibus (GEO) under accession number XXX [to be provided prior to publication].

## Results

### Specific loss of host short-form Ron protects from breast cancer bone metastasis

To investigate the role of host SF-Ron in supporting outgrowth of breast cancer bone metastasis, MMTV-PyMT mammary tumor cells were injected into the tibiae of syngeneic WT mice or mice lacking SF-Ron (Ron SF^-/-^). Two weeks after injection, MMTV-PyMT tumor cells had little effect on the bone, as assessed by X-ray imaging (**Fig. 1A** and **B**). However, by four weeks post-injection, the tumor cells caused significantly more osteolysis in WT mice than in Ron SF^-/-^ mice (**Fig. 1A** and **B**). To determine whether these differences in osteolysis were associated with altered tumor burden, we performed IHC for EpCAM, a tumor marker. While tumor burden was equivalent in WT and Ron SF^-/-^ hosts two weeks post-injection, at the four-week timepoint, the tumors were absent from most of the Ron SF^-/-^ mice, while they filled the bone marrow cavity of WT mice (**Fig. 1C** and **D**; Supplementary Fig. S1A). These findings indicated that mice lacking host SF-Ron were protected from bone metastasis outgrowth, leading us to investigate whether tumor clearance resulted from an anti-tumor immune response.

**Figure 1.**
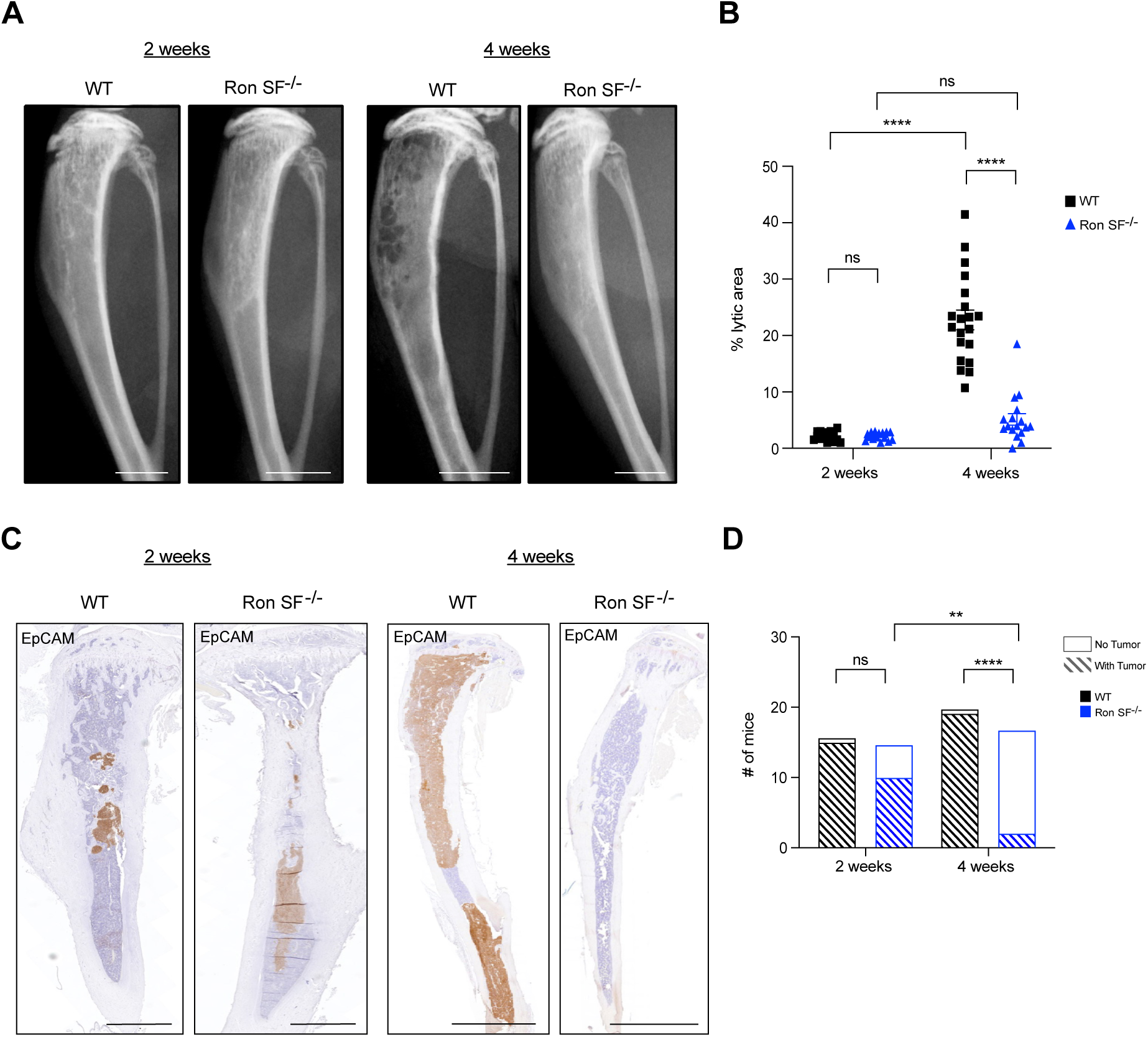
Loss of host short-form Ron promotes tumor clearance in a preclinical model of breast cancer bone metastasis. **A**, Representative X-rays of WT and Ron SF^-/-^ tibiae at 2 weeks and 4 weeks post-IT injection. Scale bars, 2 mm **B**, Lytic area quantification from WT and Ron SF^-/-^ bones at 2-week and 4-week timepoints. **C**, Representative EpCAM IHC staining of WT and Ron SF^-/-^ bones 2 weeks and 4 weeks post-IT injection. Scale bars, 1 mm at 2 weeks and 2 mm at 4 weeks. **D**, The number of mice with or without tumors, as detected by EpCAM IHC. Data are represented as mean ± SEM. *P* values were calculated using two-way ANOVA (**B**) and Fisher’s exact test (**D**). ns = not significant, *, *P* < 0.05; **, *P* < 0.01; ***, *P* < 0.001; ****, *P* < 0.0001; *n* = 16 WT, 15 Ron SF^-/-^ for 2-week timepoint and *n* = 20 WT, 17 Ron SF^-/-^ for 4-week timepoint.

### Loss of host SF-Ron enhances infiltration of immune cells into bone micrometastases

To determine whether elimination of bone micrometastases in Ron SF^-/-^ mice was related to a role for SF-Ron in regulating immune responses, we isolated bone marrow cells from WT and Ron SF^-/-^ bones during the micrometastasis stage (two weeks after IT injection) and performed flow cytometry (Supplementary Fig. S2A). Despite equivalent tumor burden at this stage, Ron SF^-/-^ bones had significantly higher numbers of CD3e+ T cells, B220+CD19+ B cells, CD11c+ dendritic cells, CD11b+F4/80+ macrophages, NK1.1+ NK cells, and CD11b+Ly6G+ neutrophils (**Fig. 2A-B),** though the relative proportions of immune cells were not consistently increased (Supplementary Fig. S2B). Interestingly, in tumor-naïve mice, we observed a significant reduction in the number of myeloid cells in the bone marrow of Ron SF^-/-^ compared with WT mice, whereas other immune cell populations were not significantly different (Supplementary Fig. S2C-D). These data suggested that Ron SF^-/-^ hosts have a more robust immune response to tumor cells in the bone.

**Figure 2.**
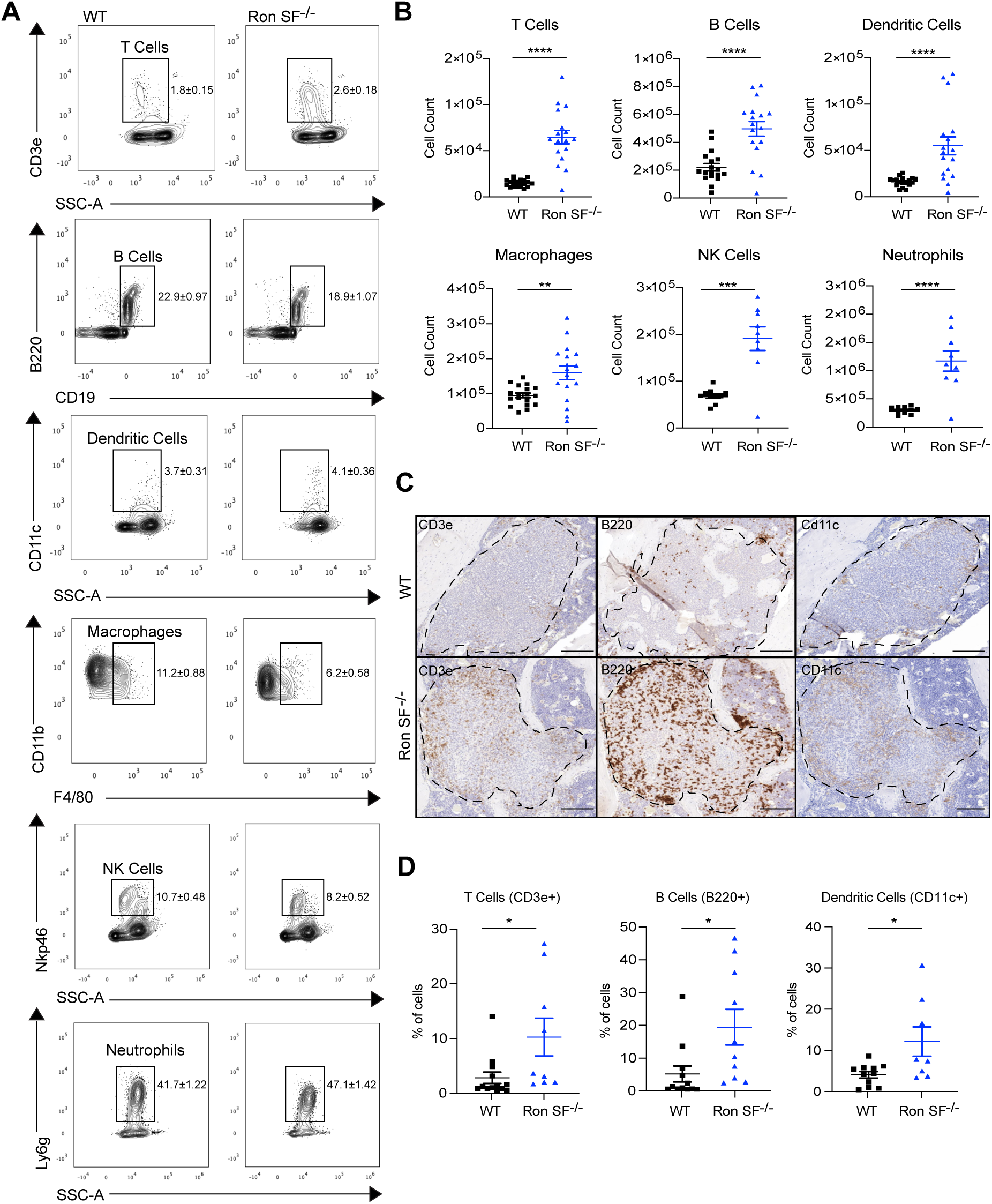
Ron SF^-/-^ tumor-bearing bones have increased immune cells and immune infiltration into tumors. **A**, Representative flow cytometry plots and gating strategy of different immune cell subsets in the bone with frequency mean ± SEM. **B**, Immune cell counts in WT (*n* = 10 or 18 per cohort) and Ron SF^-/-^ (*n* = 9 or 17 per cohort) bones, 2 weeks post-IT injection. Immune cells were defined as follows: T cells (CD3e+), B cells (B220+CD19+), dendritic cells (CD11c+), macrophages (CD11b+F4/80+), NK cells (NKp46+), neutrophils (Ly6g+). **C**, Representative IHC images of tumor-infiltrating CD3e+ (T cells), B220+ (B cells), and CD11c+ (dendritic cells) cells. Tumors are outlined. Scale bars, 100 µm. **D**, IHC quantification of tumor-infiltrating CD3e+, B220+, and CD11c+ cells from the bones of WT (*n* = 11-13) and Ron SF^-/-^ (*n* = 8-10) mice at the 2-week timepoint. Data are represented as mean ± SEM. *P* values were calculated using an unpaired two-tailed Student *t* test (**B** and **D**). *, *P* < 0.05; **, *P* < 0.01; ***, *P* < 0.001; ****, *P* < 0.0001.

Next, we asked whether specific immune cell populations had infiltrated the tumors in WT and Ron SF^-/-^ mice. Given their critical role in activating adaptive immunity for tumor clearance, we performed IHC staining for T cells, B cells, and dendritic cells. This revealed significantly more infiltrating CD3e+ T cells, B220+ B cells, and CD11c+ dendritic cells within micrometastases of Ron SF^-/-^ mice compared to WT (**Fig. 2C** and **D**). Thus, loss of host SF-Ron resulted in a robust anti-tumor immune response in the bone that also promoted immune infiltration into the micrometastases. Among the cell types we investigated, B220+ cells were the most prominent immune cell population infiltrating the tumors of Ron SF^-/-^ mice. The role of B cells in anti-tumor immunity and metastatic outgrowth is poorly understood, with few studies in bone metastasis, prompting us to investigate the B220+ population further.

### B220+ B cells exhibit inflammatory and proliferative signatures in tumor-bearing bones of Ron SF^-/-^ mice

B220 can be expressed by several immune cell types, including B cells, NK cells, T cells, and plasmacytoid dendritic cells (pDCs). Using cell depletion techniques, we determined that NK cells and pDCs were not required for Ron SF^-/-^ hosts to clear bone metastases (Supplementary Fig. S3A-D). To further determine the identity of the B220+ cells in our model, and to see if there were specific differences in B220+ cell phenotypes in the bone micrometastases of Ron SF^-/-^ hosts compared to WT, we macro-dissected tumor-enriched areas of bone marrow at the 2-week timepoint, FACS sorted the B220+ cells, and performed bulk RNA sequencing. B220+ cells from both WT and Ron SF^-/-^ hosts expressed elevated B cell signature genes in comparison to genes expressed by other B220+ cells, such as T cells, NK cells, and pDCs, which confirmed that the B220+ cells we isolated are enriched in B cells (**Fig. 3A**). Unsupervised clustering revealed strong, specific signatures in the B220+ population that were dependent on the absence of SF-Ron (**Fig. 3B**). Ron SF^-/-^ B220+ cells significantly upregulated 709 genes and downregulated 1595 genes (**Fig. 3C).** We observed significant upregulation of genes associated with B-cell activation (*Cd40*, *Cd38*, *Irf4*, *Il21r*, and *Tlr9*), antigen presentation (*Ciita*, *Cd74*, *H2-Aa*, *H2-Ab1*, and *H2-DMa*), and cell survival (*Bcl2*, *Atm*, and *Bach2*) in B220+ cells from Ron SF^-/-^ mice compared to WT mice (**Fig. 3D**). Overall, Ron SF^-/-^ B220+ cells were significantly enriched in inflammatory and proliferation gene sets, while WT B220+ cells were enriched in metabolic and tissue repair pathways (42–45) (**Fig. 3E**). We also observed that B220+ cells from Ron SF^-/-^ bones were enriched in gene expression related to B cell function, including immunoglobulin production, response to type II interferon, and antigen receptor-mediated signaling (**Fig. 3F**). Collectively, these results suggested that B cells within bone micrometastases in Ron SF^-/-^ mice have a more activated and functional transcriptional program compared to B cells isolated from bone micrometastases in WT mice.

**Figure 3.**
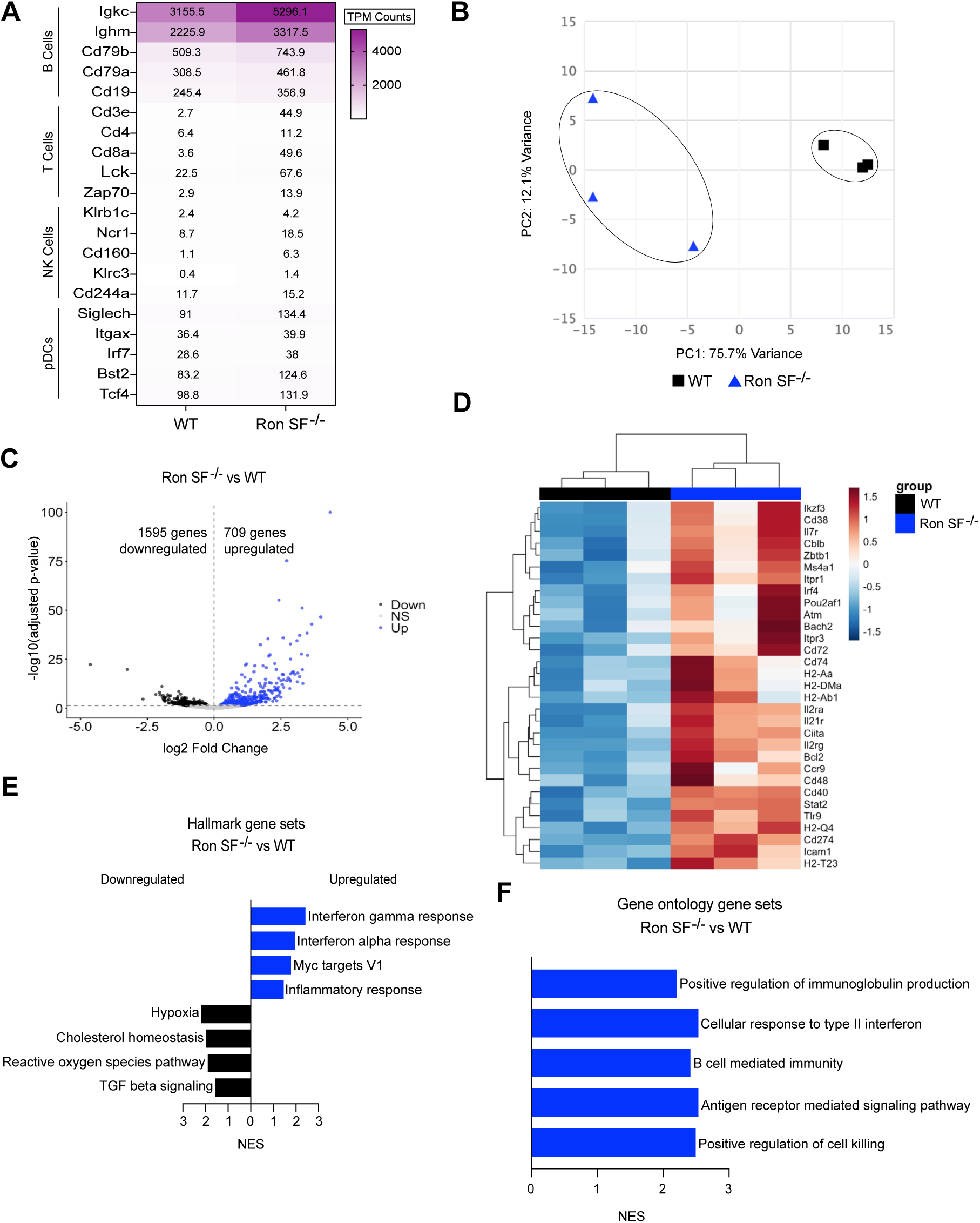
B cells from Ron SF^-/-^ hosts acquire an active and inflammatory transcriptional state in the presence of tumors. **A,** Average transcripts per million (TPM) counts for select genes of B cell, T cell, NK cell, and pDC populations. **B**, PCA map based on RNA sequencing of FACS-isolated B220+ cells from WT (black, *n* = 3) and Ron SF^-/-^ (blue, *n* = 3) tumor-bearing mice. **C**, Volcano plot showing DEGs [-log10 (*P*adj) > 1.3] upregulated (right, blue) or downregulated (left, back) in Ron SF^-/-^ vs WT B220+ cells. **D**, Heatmap showing differentially expressed B-cell-associated genes (*P*adj < 0.05). The color scale represents the normalized Z score between Ron SF^-/-^ and WT samples. **E**, Bar plots showing select significantly upregulated (blue) and downregulated (black) Hallmark gene sets in Ron SF^-/-^ B cells by GSEA. **F**, Upregulated Gene Ontology gene sets in Ron SF^-/-^ B cells by GSEA. For GSEA, *P* value < 0.05, FDR <0.25, and NES <-1 or >1 were considered significant. Genes were preranked based on log_2_ fold change (**E** and **F**).

### Mature B cells are required for protection from bone metastasis in Ron SF^-/-^ mice

To definitively determine whether B cells were required to prevent bone metastasis in Ron SF^-/-^ mice, we genetically depleted B cells by crossing Ron SF^-/-^ mice to JH^-/-^ mice, which lack the gene encoding the heavy chain joining region and cannot produce functional B cells (34). We confirmed that double knockout mice (Ron SF^-/-^JH^-/-^) have significantly reduced B cells (**Fig. 4A**). Four weeks post-intratibial injection of MMTV-PyMT tumor cells, we observed osteolysis consistent with tumor growth in WT and JH^-/-^ mice, while Ron SF^-/-^ mice had little osteolysis, as expected. In contrast, Ron SF^-/-^JH^-/-^ mice had significantly increased osteolysis compared to the Ron SF^-/-^ cohort, suggesting bone metastasis had been rescued in Ron SF^-/-^ mice lacking functional B cells (**Fig. 4B**). Indeed, outgrowth of metastases in Ron SF^-/-^JH^-/-^ bones was confirmed by EpCAM staining at the four-week timepoint, where a significantly greater proportion of Ron SF^-/-^JH^-/-^ hosts developed tumors compared to Ron SF^-/-^ hosts. We observed no differences between WT and JH^-/-^ mice, both of which developed tumors (**Fig. 4C**). As expected, the B cell infiltrative phenotype of Ron SF^-/-^ mice was lost in Ron SF^-/-^JH^-/-^ hosts (**Fig. 4D** and **E**). Taken together, these data demonstrated not only that loss of SF-Ron promoted B cell infiltration and activation in bone micrometastases, but also that these B cells contributed to tumor elimination: functional B cells were required for protection from bone tumors in Ron SF^-/-^ hosts.

**Figure 4.**
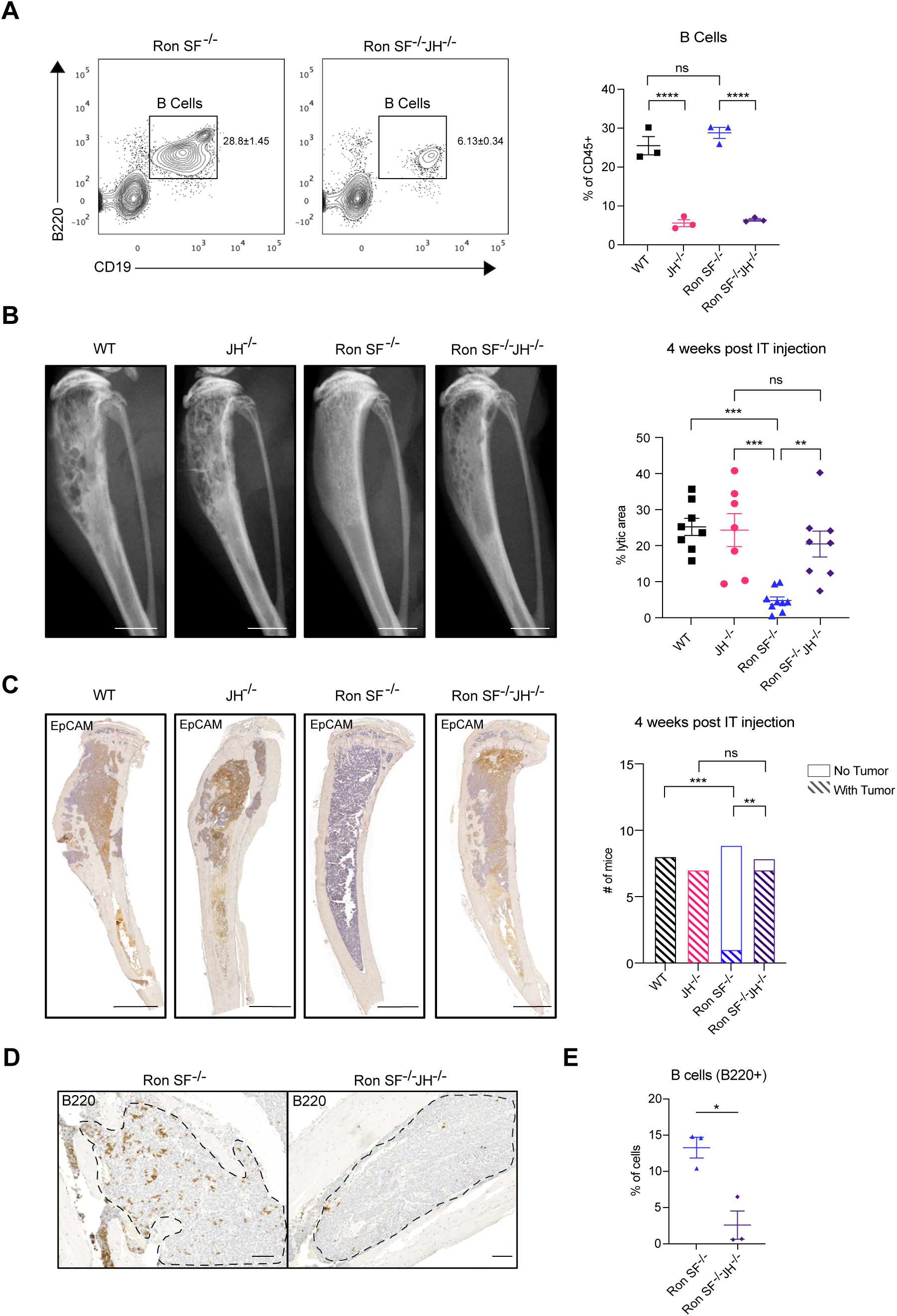
Loss of B cells rescues tumor growth in Ron SF^-/-^ mice. **A**, Representative flow plot (left) and quantification (right) of B220+CD19+ bone marrow B cell frequency in Ron SF^-/-^ and Ron SF^-/-^ JH^-/-^ mice (*n* = 3/group). **B**, Representative X-rays (left) and lytic area quantification (right) of WT, JH^-/-^, Ron SF^-/-^, and Ron SF^-/-^JH^-/-^ bones, 4 weeks post-IT injection (*n* = 7-9/group). Scale bars, 2 mm **C**, Representative EpCAM IHC images (left) and quantification of the number of mice with tumors (right) of WT, JH^-/-^, Ron SF^-/-^, and Ron SF^-/-^JH^-/-^ bones, 4 weeks post-IT injection. Scale bars, 1 mm. **D**, Representative IHC images of tumor-infiltrating B220+ cells (B cells) at the 2-week timepoint in Ron SF^-/-^ and Ron SF^-/-^JH^-/-^ bones. Scale bars, 100 µm. **E**, Quantification of tumor-infiltrating B220+ cells at the 2-week timepoint in Ron SF^-/-^ (*n* = 3) and Ron SF^-/-^JH^-/-^ (*n* = 3) mice. Tumors are outlined. Data are represented as mean ± SEM. *P* values were determined using one-way ANOVA (**A** and **B**), Fisher’s exact test (**C**), and unpaired two-tailed Student *t* test (**E**). ns = not significant, *, *P* < 0.05; **, *P* < 0.01; ***, *P* < 0.001; ****, *P* < 0.0001.

### Mature B cells in Ron SF^-/-^ mice cooperate with T cells to eliminate tumors in the bone

To better understand why B cells from Ron SF^-/-^ mice were critical for effectively clearing bone micrometastases, we performed single-cell RNA sequencing (scRNA-seq). We isolated B220+ cells by FACS from tumor-enriched regions of WT and Ron SF^-/-^ hosts at the two-week time point and analyzed 22,095 cells from 10 different mice (5 WT and 5 Ron SF^-/-^). After excluding B220+ immune cells that do not express the B cell-associated genes *Cd19*, *Cd79a*, *Cd79b*, and *Ms4a1,* unsupervised clustering revealed 7 clusters of B cell subsets (**Fig. 5A**). The proportion of each cluster was not significantly different between genotypes (Supplementary Fig. S4A-B). We annotated B cell subsets using genes associated with published B cell subsets and the web-based Cluster Identity Predictor (CIPR) tool, which compares cluster expression signatures with those of defined immune cell populations from the Immunological Genome Project database (41) (**Fig. 5B**; Supplementary Fig. S4C). The clusters detected were follicular B cells, transitional B cells, two pre-B cell populations, cycling pre-B cells, pre-pro-B cells, and pro-B cells (**Fig. 5B**). Most of the clusters were associated with developing B cells, which are abundant in the bone marrow. We focused our subsequent analysis on the mature B cell clusters because JH^-/-^ mice only lack mature B cells (34), functionally implicating them in protection against metastatic outgrowth in Ron SF^-/-^ mice. Clusters 2 and 6 were enriched for genes expressed by mature B cells, including *Cd19*, *Ms4a1*, *Ighd,* and *Cd40* (**Fig. 5B**). Cluster 2 was associated with a transitional B cell signature using CIPR. Cluster 6 was enriched in follicular B cell genes such as *Fcer2a*, *Cr2*, and *Sell,* and was associated with a follicular B cell transcriptional program through CIPR (**Fig. 5B**). Further investigation into each cluster revealed that transitional B cells from Ron SF^-/-^ mice were enriched in proliferative and inflammatory gene sets (**Fig. 5C-E**), while follicular B cells displayed upregulated active and inflammatory programs and interferon gamma response genes (**Fig. 5F-H**). We found no significant enrichment of pathways in mature B cell subsets from WT bones.

**Figure 5.**
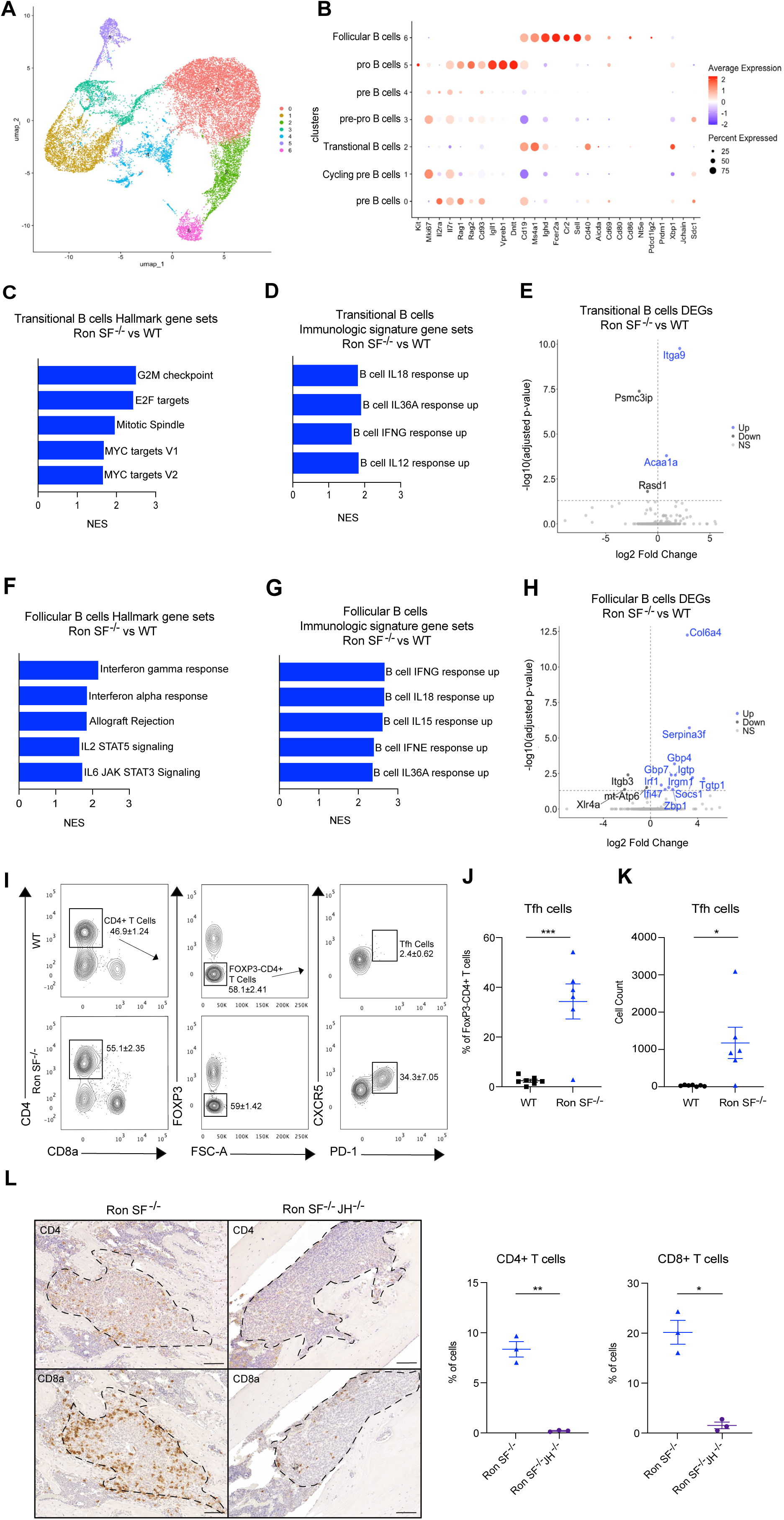
Transcriptional profiling for specific B cell subsets reveals enrichment for transitional B cells and follicular B cells in Ron SF^-/-^ tumor-bearing hosts. **A**, UMAP projection of scRNA seq data showing different clusters of B220+ cells from tumor-bearing Ron SF^-/-^ (*n* = 5) and WT (*n*=5) mice at the 2-week timepoint. **B**, Dot plot showing expression of representative genes of each B cell cluster, annotated with CIPR. Dot size reflects the percentage of cells in a cluster expressing the gene; dot colors indicate average expression levels. **C**, Upregulated Hallmark gene sets by GSEA in Ron SF^-/-^ transitional B cells. **D**, Upregulated selected pathways by GSEA using immunologic signature gene sets in transitional B cells from Ron SF^-/-^ mice. **E**, Volcano plot of DEGs [-log10 (*P*adj) > 1.3] upregulated (right, blue) or downregulated (left, black) in Ron SF^-/-^ vs WT transitional B cells. **F**, Upregulated selected pathways by GSEA using Hallmark gene sets in follicular B cells from Ron SF^-/-^ mice. **G,** Upregulated selected pathways by GSEA using immunologic signature gene sets in follicular B cells from Ron SF^-/-^ mice. **H**, Volcano plot showing DEGs [-log10 (*P*adj) > 1.3] upregulated (right, red) or downregulated (left, blue) in Ron SF^-/-^ vs WT follicular B cells. **I**, Representative flow cytometry plots of the gating strategy for Tfh cells in the bone, showing frequency as mean ± SEM. **J**, Frequency of Tfh cells in the bones of WT (*n* = 7) and Ron SF^-/-^ (*n* = 6) mice, 2 weeks post-IT injection. **K,** Cell counts of Tfh cells in the bones of WT (*n* = 7) and Ron SF^-/-^ (*n* = 6) mice, 2 weeks post-IT injection. **L,** Representative IHC images (left) and quantification (right) of tumor-infiltrating CD4+ and CD8+ T cells in Ron SF^-/-^ and Ron SF^-/-^JH^-/-^ bones at the two-week timepoint. Tumors are outlined. Scale bars, 200 µm. Data are represented as mean ± SEM. For GSEA, *P* value < 0.05, FDR <0.25, and NES <-1 or >1 were considered significant. Genes were preranked based on log_2_ fold change (**D-H**).

We noted enrichment of cytokine response programs in mature B-cell subsets, as well as the presence of transcriptionally inflammatory follicular B cells in Ron SF^-/-^ bones (**Fig. 5D** and **G-H**), suggesting that crosstalk between B cells and other immune cells may contribute to the elimination of micrometastases. Through antigen presentation and cytokine production, follicular B cells work extensively with T follicular helper (Tfh) cells to mount immune responses (46,47). Accordingly, we detected a large, significant increase in the proportion and number of Tfh cells in the tumor-enriched areas of Ron SF^-/-^ bones relative to WT bones by flow cytometry **(Fig. 5I-K).** This increase in Tfh cells was driven by the presence of tumors, as we observed no evidence of Tfh cells in tumor-naïve bones (Supplementary Fig. S4D and E).

As previously noted, micrometastases in Ron SF^-/-^ mice were infiltrated with both B cells and T cells (**Fig. 3E**). To determine if the B cells were responsible for recruiting T cells into tumors, we asked if elimination of mature B cells in Ron SF^-/-^ hosts (Ron SF^-/-^JH^-/-^) altered T cell infiltration into the bone micrometastases. IHC staining for CD4 and CD8 revealed that infiltration of both CD4+ and CD8+ T cells into micrometastases was absent when mature B cells were depleted from Ron SF^-/-^ mice (**Fig. 5L**).

Finally, to investigate the necessity of T cells in the clearance of bone metastases in SF^-/-^ mice, we depleted CD4+ or CD8+ T cells by crossing Ron SF^-/-^ mice to CD4^-/-^ or CD8^-/-^ mice, respectively (Supplementary Fig. S5A-B). Double knockouts had significantly more osteolytic lesions than Ron SF^-/-^ controls when injected with MMTV-PyMT tumor cells (Supplementary Fig. S5C-D). IHC staining four weeks after injection confirmed that the loss of either CD4+ or CD8+ T cells in Ron SF^-/-^ mice resulted in significantly more bone metastasis, rescuing the Ron SF^-/-^ phenotype (**Fig. 6A-B**). Interestingly, we also observed a reduction in B-cell tumor infiltration when CD4+ or CD8+ T cells were depleted in Ron SF^-/-^ hosts (**Fig. 6C**), demonstrating that a virtuous cycle of tumor infiltration by both B and T cells is promoted by depletion of host SF-Ron. Together, these data show that distinct B cell subsets, including follicular B cells, are specifically recruited into tumors and activated in Ron SF^-/-^ hosts, resulting in an ensuing Tfh response and the recruitment of both CD4+ and CD8+ T cells into micrometastases.

**Figure 6.**
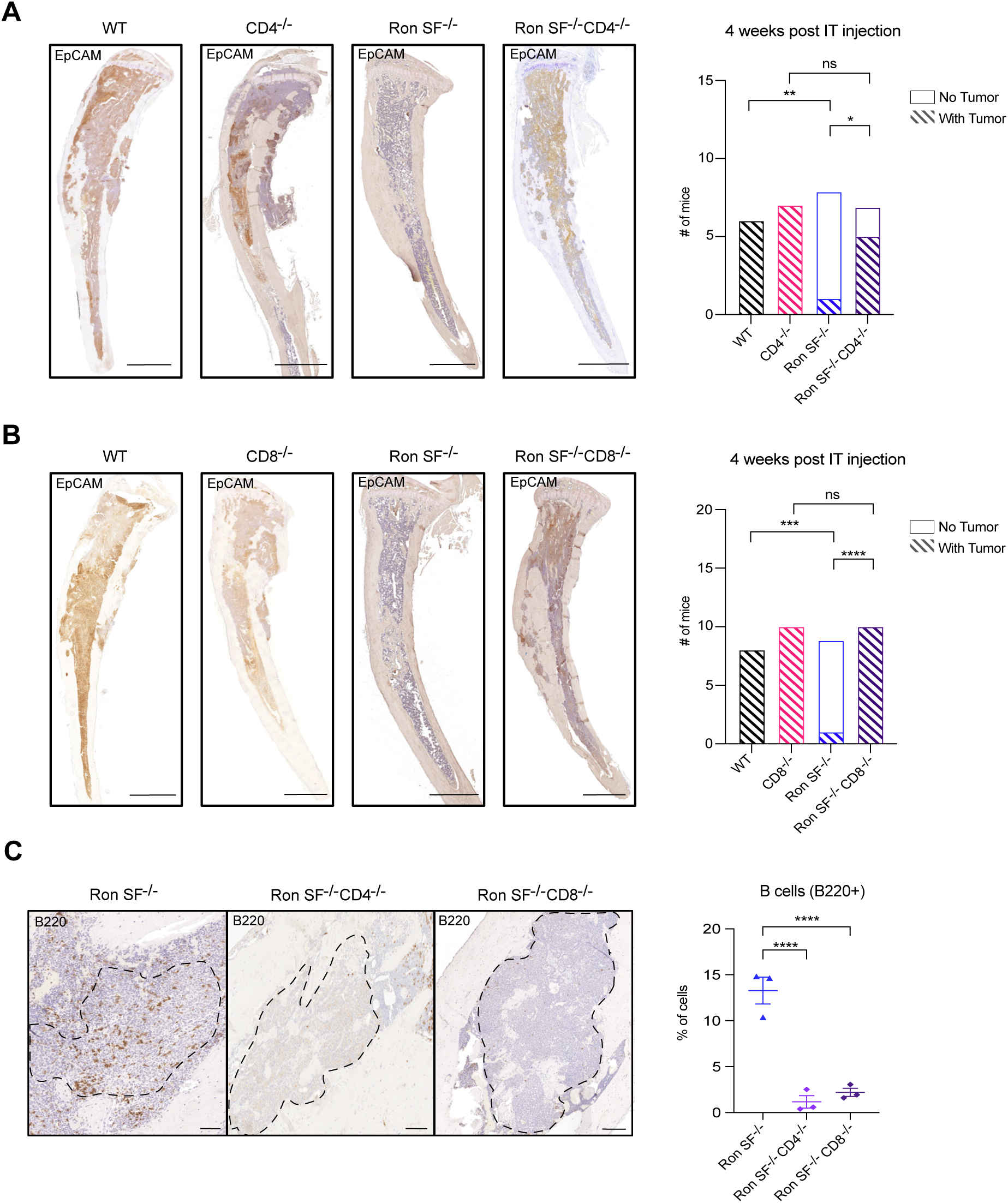
T cells are necessary for SF-Ron-mediated protection against tumor growth in the bone. **A**, Representative images of EpCAM IHC staining of WT, CD4^-/-^, Ron SF^-/-^, and Ron SF^-/-^CD4^-/-^ bones (left), with quantification of the number of mice with tumors (right) at the 4-week timepoint. Scale bars, 1 mm. **B**, Representative images of EpCAM IHC staining of WT, CD8^-/-^, Ron SF^-/-^, and Ron SF^-/-^CD8^-/-^ bones (left) with quantification of tumors (right), 4 weeks post-IT injection, Scale bars, 1 mm. **C**, Representative IHC images of tumor-infiltrating B220+ (B cells) (left) and quantification (right) at the 2-week timepoint in Ron SF^-/-^, Ron SF^-/-^CD4^-/-^, and Ron SF^-/-^CD8^-/-^ bones. Scale bars, 100 µm. Data are represented as mean ± SEM. *P* values were calculated using Fisher’s exact test (**A** and **B**) and an unpaired two-tailed Student *t* test (**C**). ns = not significant, *, *P* < 0.05; **, *P* < 0.01; ***, *P* < 0.001; ****, *P* < 0.0001.

### B and T cell cooperation is required for the prevention of metastatic outgrowth in bones, but not lungs, of Ron SF^-/-^ mice

The necessity of T cells in SF-Ron-mediated protection against bone metastasis parallels our previously published work identifying a role for T cells in SF-Ron-mediated protection against lung metastasis (31). Therefore, we asked whether mature B cells are also required for protection from lung metastasis, as observed in the bone. Similar to the bone, a greater proportion of Ron SF^-/-^ mice are protected from lung metastasis compared to WT (**Fig. 7A**). However, it was remarkable that mature B cells were not required to prevent lung metastasis, as the majority of Ron SF^-/-^JH^-/-^ mice remained protected (**Fig. 7A** and **B** and Supplementary Fig. S6). In both Ron SF^-/-^ and Ron SF^-/-^JH^-/-^ mice, we noted instances of mice with tumors that escaped immune clearance and had similar tumor burden to WT (**Fig. 7B** and Supplementary Fig. S6). This phenomenon is also occasionally observed in the bone, indicating that tumor control in Ron SF^-/-^ is effective in most, but not all, hosts. Nevertheless, these findings uncover a clear role for SF-Ron in controlling anti-tumor immunity against metastasis, with mature B cells being essential for immune protection against bone but not lung metastasis in the absence of SF-Ron.

**Figure 7.**
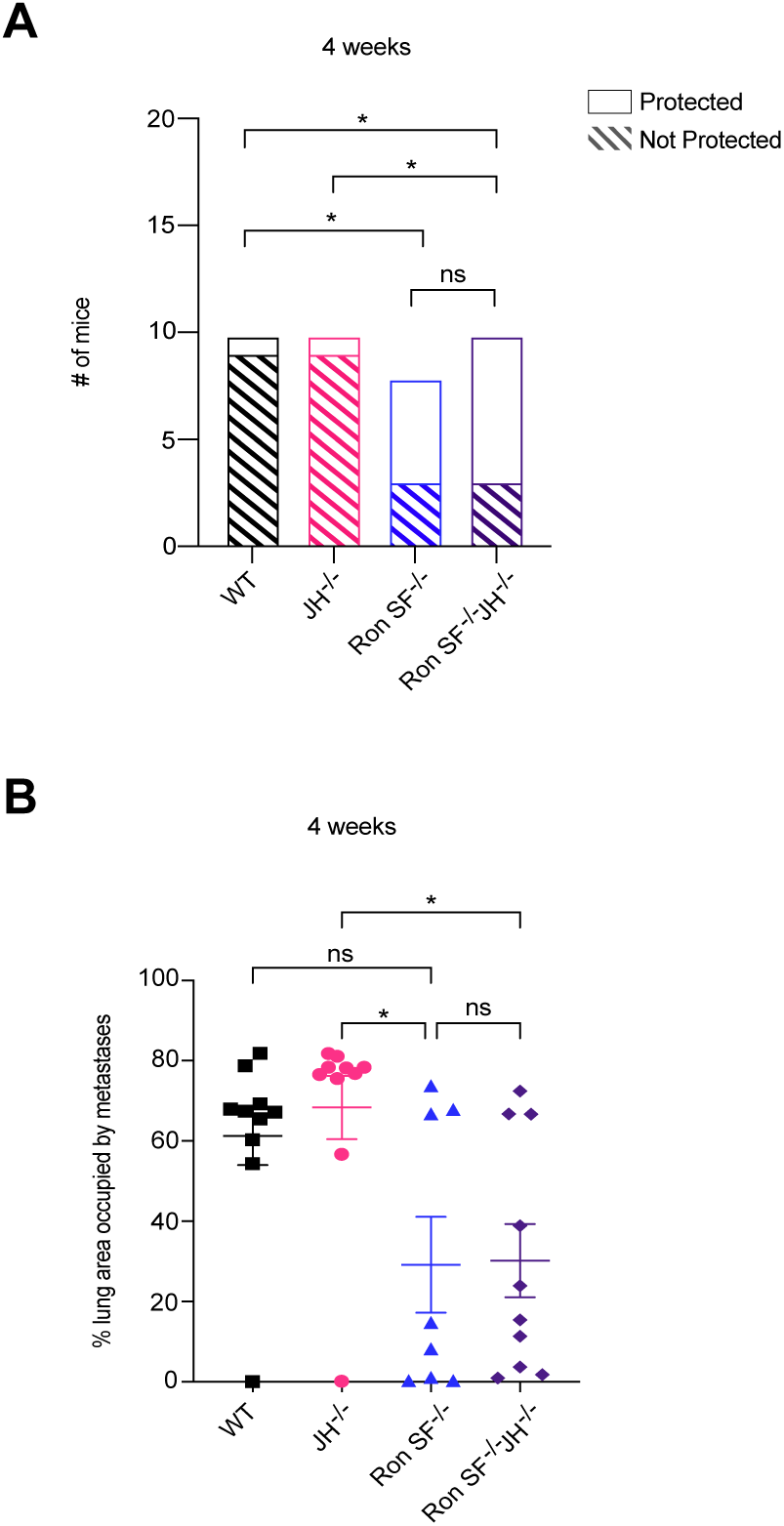
B cells are not required for SF-Ron-mediated protection against metastasis outgrowth in the lung. **A**, The number of mice protected or not from tumor growth 4 weeks post-TV injection. **B**, Quantification of the percent lung area occupied by metastasis (n = 8–10 mice/group). Data are represented as mean ± SEM. *P* values were determined using Fisher’s exact test (**A**) and one-way ANOVA (**B**). ns = not significant, *, *P* < 0.05.

## Discussion

This study establishes, for the first time, a specific role for host SF-Ron in breast cancer bone metastasis. We show that micrometastatic tumors establish similarly in the bones of WT mice and mice lacking SF-Ron, but are later cleared in Ron SF^-/-^ hosts in a B cell- and T cell-dependent manner. Notably, B cells are not required for efficient anti-tumor immunity in a lung metastasis model. Our data provide compelling evidence that host short-form Ron plays a previously unknown role in organ-dependent B cell–T cell cooperation, whereby loss of SF-Ron promotes strong functional cooperation between B cells and T cells, sufficient to eliminate micrometastases and prevent bone metastatic outgrowth. While these data should be confirmed in additional models, including spontaneous bone metastasis models, when taken together with our previous report (31), the collective data strongly implicate a major role for SF-Ron in regulating anti-tumor immunity across multiple metastatic sites, including the lung and bone. Although bone-only metastasis is not itself terminal, it causes major skeletal-related complications (6,7), is implicated in promoting the spread of cancer cells to other tissues (2,48), and can influence systemic responses to immunotherapy (15,16). Thus, enhancing anti-tumor immunity against bone metastasis may positively impact patient outcomes beyond the local effects in the bone. Our findings warrant focusing on targeting SF-Ron to prevent metastatic progression. The non-redundant roles of FL-Ron and SF-Ron in cancer and inflammatory conditions (21,30) support the development of therapies targeting SF-Ron and preclinical studies to evaluate their efficacy in combination with approved cancer treatments, such as immunotherapy.

In the field of anti-tumor immunity, T cells have been the primary focus, while B cells have been understudied. However, recent findings suggest a need and interest to better understand the role of B cells in anti-tumor immunity (49,50). In animal models, B cells have been implicated in promoting tumor growth as immunosuppressive regulatory B cells (Bregs), and in disrupting the anti-tumor function of T cells through modulation of regulatory CD4+ T cells (51). Conversely, in breast cancer patients, B cell infiltration into tumors has been associated with improved outcomes (52,53). Little is known about the role of B cells in metastatic breast cancer, but a few studies have shown their pro- and anti-tumor capabilities in breast cancer lung metastasis models (51,54). To our knowledge, this study is the first to demonstrate that B cells are necessary for anti-tumor immune response in bone metastasis. B cells were the predominant immune cells infiltrating bone micrometastases in Ron SF^-/-^ mice, suggesting that targeting host SF-Ron could improve outcomes for metastatic breast cancer patients, in part by stimulating B cell infiltration and function in bone metastases.

The anti-tumor function of B cells has been primarily attributed to the formation of tertiary lymphoid structures, which are associated with improved patient outcomes (52,55). B cells also contribute to anti-tumor immunity through antigen presentation and antibody production (56,57). Our bulk and single-cell RNA sequencing revealed that mature B cells from tumor-bearing Ron SF^-/-^ bones were more active and inflammatory than WT B cells. Ron SF^-/-^ B cells upregulated antigen-presentation genes, such as *Ciita* and *Cd74*, and genes associated with T cell crosstalk, like *Cd40* and *Il21r*. Furthermore, Ron SF^-/-^ follicular B cells exhibited increased expression of interferon-responsive genes, implying that cooperation between B cells and other immune cells may promote the clearance of micrometastases. Others have shown that B cell-dependent anti-tumor immune responses are frequently associated with Tfh cells (58,59). In breast cancer mouse models, B cells and Tfh cells were necessary for effective anti-tumor immune responses with immune checkpoint blockade in highly immunogenic triple-negative primary mammary tumors (58). This phenomenon was similarly observed in a primary lung cancer model, in which B cells were required to induce Tfh cells and promote their IL-21 production and CD8+ T cell activation (59). Here, we found that the tumor microenvironment of Ron SF^-/-^ bones was characterized by both follicular B cells and abundant Tfh cells. These findings are the first to implicate SF-Ron in B cell function and suggest a previously undescribed B cell-Tfh axis in bone metastasis. Future studies are pertinent to define the underlying role of SF-Ron in B cell function and in B and T cell cooperation that contribute to anti-tumor immunity.

It has been appreciated that tissue-specific microenvironments can profoundly influence primary tumor growth and metastatic progression, with the composition and functional status of the immune microenvironment playing an important role in controlling tumor progression (60–62). Our findings extend this concept by showing that loss of host SF-Ron protects against metastatic outgrowth through distinct tissue-dependent immune mechanisms, most notably by promoting anti-tumor B and T cell cooperation in the bone, which is not required in the lung. Similar organ-dependent immune requirements have been observed in colorectal cancer, where loss of TFGβ signaling in myeloid cells reduced metastatic outgrowth in both the lung and liver through distinct mechanisms, namely, a requirement for CD8+ T cells was found in the lung but not the liver (63). Together, these findings support the emerging concept that the effects of modulating immune-associated pathways on tumor growth can differ across tissues. Here, we identified SF-Ron as a regulator of organ-specific anti-tumor immunity, revealing a distinct role for B and T cell cooperation in controlling metastatic outgrowth in the bone. These findings underscore the importance of studying metastatic sites as distinct immune microenvironments, in which the tissue niche can determine whether an anti-tumor response occurs and the mechanisms by which tumor control is achieved.

## Acknowledgements

We thank Dr. Lisa Coussens for the generous gift of CD4^-/-,^ CD8^-/-^, and JH^-/-^ cryopreserved sperm from homozygous mice. We acknowledge the flow cytometry and cell imaging core facilities at the University of Utah for access to equipment, including the BD LSRFortessa flow cytometer and Zeiss Axioscan Z1, and thank James Marvin, Eduardo Salustiano Jesus dos Santos, Sreeja Govindarajan, and Anton Classen for their assistance with training and sample acquisition. We gratefully acknowledge support from the Department of Defense Breast Cancer Research Program (W81XWH1810616 to ALW), the Susan G. Komen Foundation (SAC220226 to ALW), the Breast Cancer Research Foundation (BCRF-24-200 to ALW), the Biorepository and Molecular Pathology and High-Throughput Genomics and Cancer Bioinformatics Shared Resources, supported by HCI’s National Cancer Institute Cancer Center Support Grant P30CA042014. We thank Drs. Anna Beaudin, Kaitlyn Basham, Matthew Williams, Kevin B. Jones, and members of the Welm lab for helpful discussions of the manuscript. We acknowledge the use of OpenAI’s ChatGPT (GPT-5) to assist with improving the clarity and readability of the manuscript. All AI-assisted content was reviewed and edited by the authors, who take full responsibility for the accuracy and integrity of the final manuscript.

## Author contributions

**C. Valencia**: Conceptualization, formal analysis, validation, investigation, visualization, methodology, writing–original draft, writing–review and editing. **M. Nadjsombati**: Conceptualization, methodology, investigation, writing–review and editing. **T. Pham:** Investigation, writing–review and editing. **N. Devarajan**: Methodology, investigation. **D. Soltero**: Methodology. **J. Fornetti**: Conceptualization, formal analysis, visualization, supervision, investigation, writing–original draft, writing–review and editing. **A. Welm**: Conceptualization, funding acquisition, resources, project administration, supervision, investigation, writing original draft, writing–review and editing.

## Synopsis

We identified short-form Ron as a key regulator of tissue-specific anti-tumor B and T cell crosstalk that regulates the development of breast cancer bone metastasis.

## Conflict of interest statement

The authors declare no conflicts of interest.

## Notes

### Competing Interest Statement

The authors have declared no competing interest.

