## Supplementary data for "Loss of short-form Ron promotes B and T cell cooperation to prevent outgrowth of breast cancer bone metastasis"

### Supplementary Materials

| Mouse alleles | Primers |
|---|---|
| WT CD8<br>305 bp | Forward 5′ - GGACCTGGTATGTGAAGTGTTGG - 3′<br>Reverse 5′ - TTTCTGAAGGACTGGCACGACAG - 3′ |
| $CD8^{-/-}$<br>385 bp | Forward 5′ - GGACCTGGTATGTGAAGTGTTGG - 3′<br>Reverse 5′ - ATGGCGATGCCTGCTTGCCGAA - 3′ |
| WT CD4<br>634 bp | Forward 5′ - GAGGTTCGCCTTCGCAGTTTGAT - 3′<br>Reverse 5′ - GGTGCAGTTCCAGAAGTCGCT - 3′ |
| $CD4^{-/-}$<br>393 bp | Forward 5′ - GAGGTTCGCCTTCGCAGTTTGAT - 3′<br>Reverse 5′ - ATGGCGATGCCTGCTTGCCGAA - 3′ |
| WT JH<br>252 bp | Forward 5′ - CCCCACCATCACAGACCTTT - 3′<br>Reverse 5′ - ACCTTGCACCAGTCAGAGAC - 3′ |
| $JH^{-/-}$<br>375 bp | Forward 5′ - GATGGATTGCACGCAGGTTCT - 3′<br>Reverse 5′ - AGGTAGCCGGATCAAGCGTAT - 3′ |

**Table. S1. Genotyping primers used for mouse models.**

The primers are listed for each mouse model used to genetically deplete CD4+ T cells, CD8+ T cells, and mature B cells. Each primer pair corresponds to either the WT or the mutant allele, with the expected product sizes.

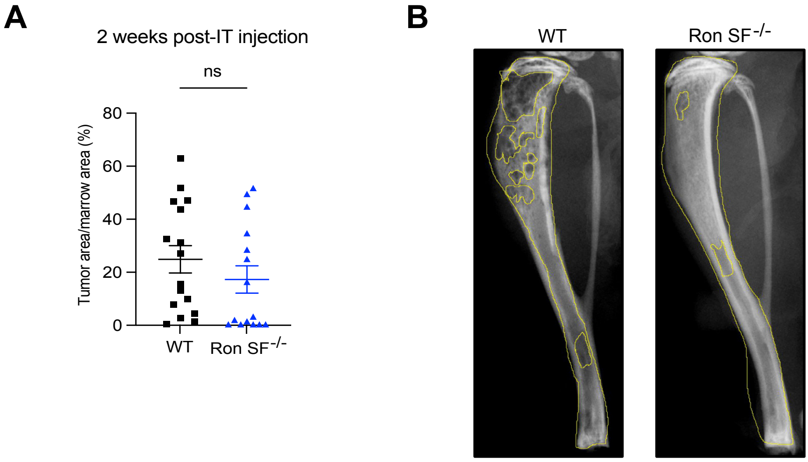

A
2 weeks post-IT injection
ns
Tumor area/marrow area (%)
80
60
40
20
0
WT
Ron SF-/-
B
WT
Ron SF-/-

**Fig. S1**. **Tumors initially grow in Ron SF$^{-/-}$ bones but are later cleared.** **A**, Quantification of the percent of bone marrow occupied by EpCAM+ tumor cells in WT and Ron SF$^{-/-}$ mice, 2 weeks post-IT injection ($n$ = 16 WT, 15 Ron SF$^{-/-}$). **B**, Representative X-rays of Ron SF$^{-/-}$ and WT tibias displaying lytic area quantification (outlined), 4-weeks post-IT injection. Data are represented as mean ± SEM. $P$ value was calculated using an unpaired two-tailed Student $t$ test. ns = not significant, $P \geq 0.05$.

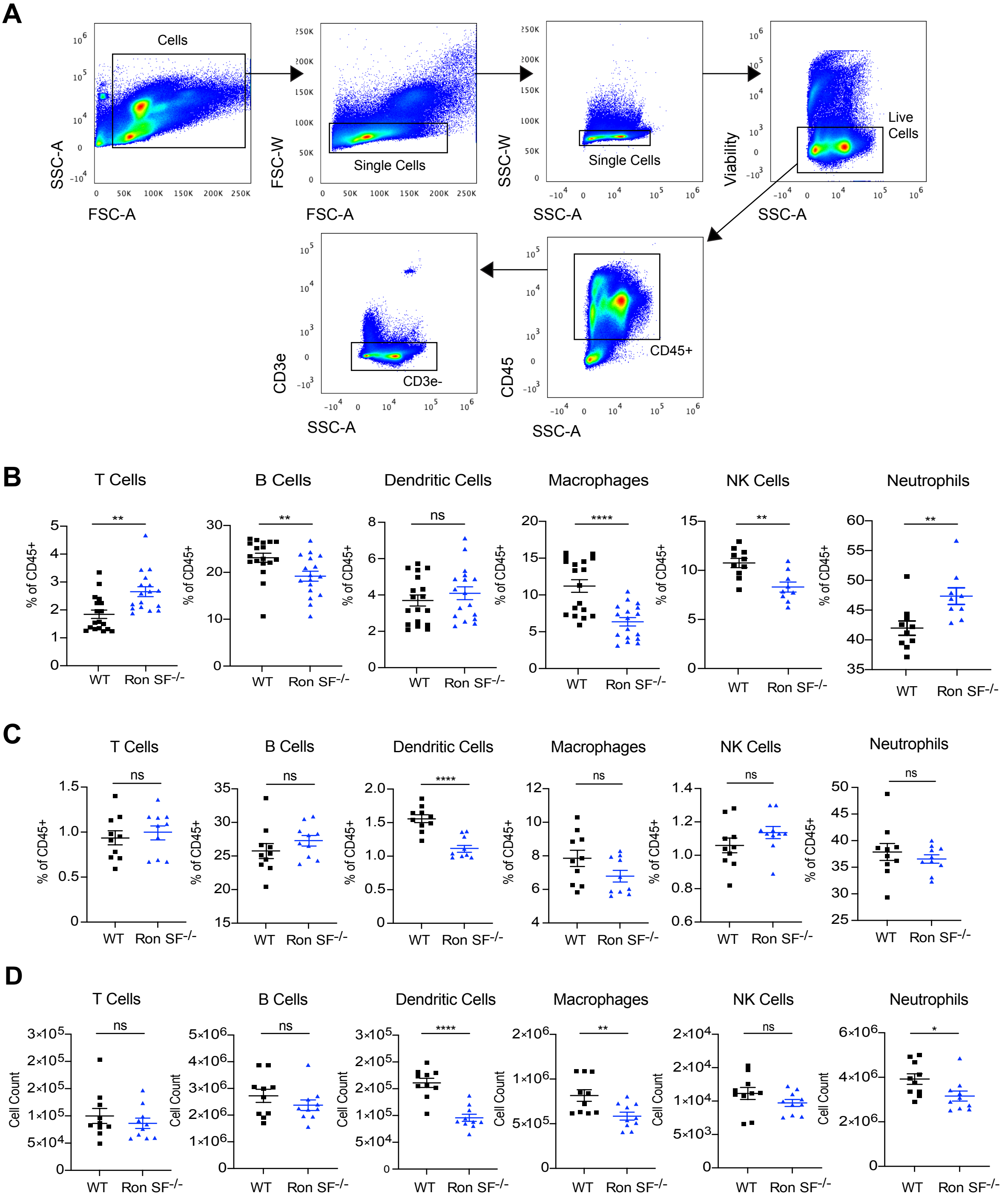
A
Cells
Single Cells
Single Cells
Live Cells
CD45+
CD3e-
SSC-A
FSC-A
FSC-W
SSC-W
Viability
CD45
CD3e
B
C
D
T Cells
B Cells
Dendritic Cells
Macrophages
NK Cells
Neutrophils
% of CD45+
Cell Count
WT
Ron SF-/-
ns
**
****
*

**Fig. S2**. **Immune cells in tumor naive bone marrow are not increased by the loss of SF-Ron.** **A**, Representative flow cytometry plots of the initial gating strategy before analyzing specific immune cell subsets. **B**, Frequency of T cells, B cells, dendritic cells, macrophages, NK cells, and neutrophils 2 weeks post-IT injections in the bones of WT ($n$ = 10-18) and Ron SF$^{-/-}$ ($n$ = 9-17). **C**, Frequency of T cells, B cells, dendritic cells, macrophages, NK cells, and neutrophils in the bone marrow of tumor naïve WT ($n$ = 10) and Ron SF$^{-/-}$ ($n$ = 10) mice. **D**, Cell numbers of T cells, B cells, dendritic cells, macrophages, NK cells, and neutrophils in the bone marrow of tumor naïve WT and Ron SF$^{-/-}$ mice. Data are represented as mean ± SEM. $P$ values were calculated using an unpaired two-tailed Student $t$ test. ns = not significant, $P \geq 0.05$; *, $P < 0.05$; **, $P < 0.01$; ***, $P < 0.001$; ****, $P < 0.0001$.

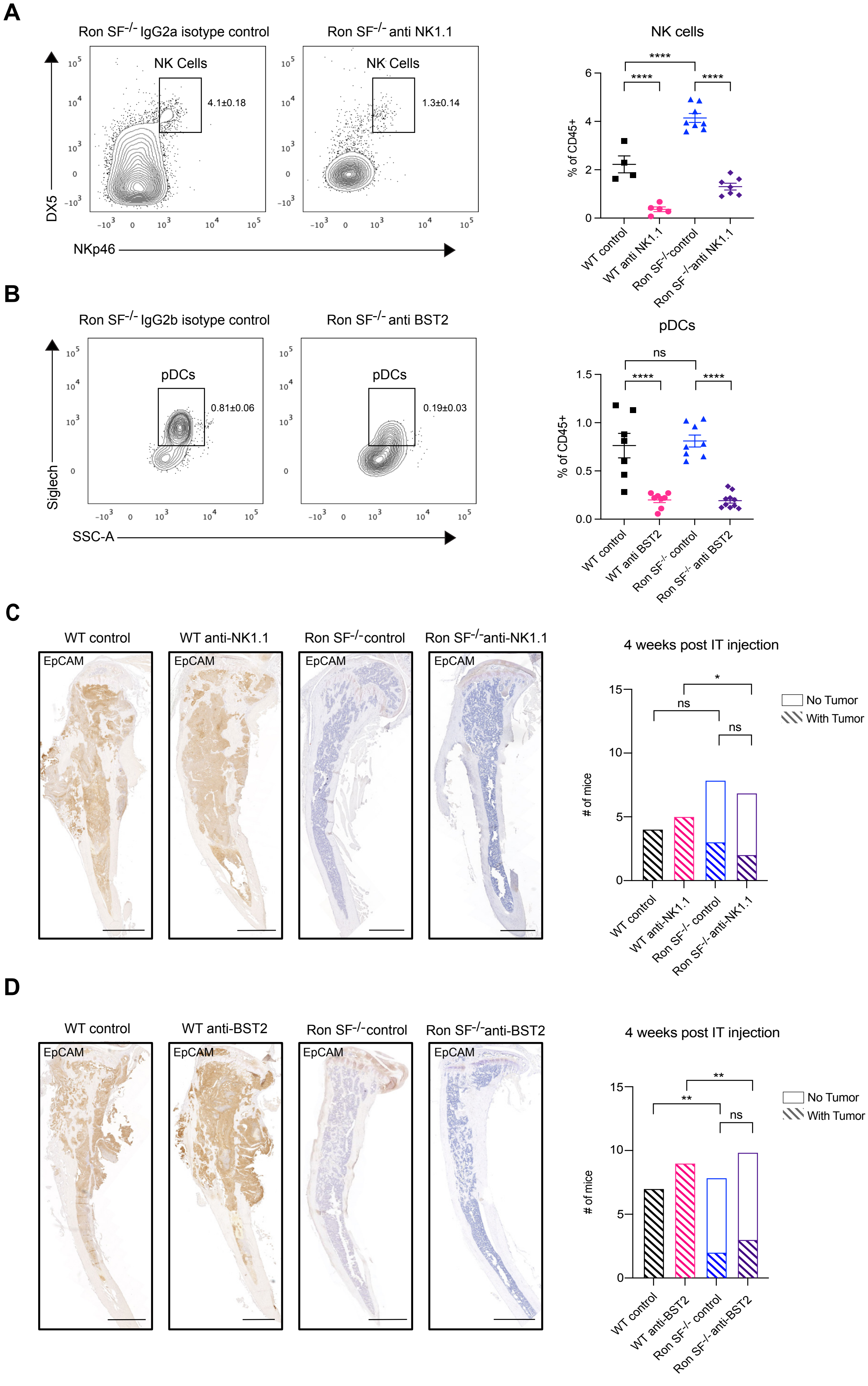
A
Ron SF-/- IgG2a isotype control
Ron SF-/- anti NK1.1
NK Cells
4.1±0.18
1.3±0.14
DX5
NKp46
NK cells
****
% of CD45+
WT control
WT anti NK1.1
Ron SF-/-control
Ron SF-/-anti NK1.1
B
Ron SF-/- IgG2b isotype control
Ron SF-/- anti BST2
pDCs
0.81±0.06
0.19±0.03
Siglech
SSC-A
ns
% of CD45+
WT control
WT anti BST2
Ron SF-/- control
Ron SF-/- anti BST2
C
WT control
WT anti-NK1.1
Ron SF-/-control
Ron SF-/-anti-NK1.1
EpCAM
4 weeks post IT injection
*
ns
No Tumor
With Tumor
### of mice
WT control
WT anti-NK1.1
Ron SF-/- control
Ron SF-/-anti-NK1.1
D
WT control
WT anti-BST2
Ron SF-/-control
Ron SF-/-anti-BST2
EpCAM
4 weeks post IT injection
**
ns
No Tumor
With Tumor
### of mice
WT control
WT anti-BST2
Ron SF-/- control
Ron SF-/-anti-BST2

**Fig. S3**. **NK cells and plasmacytoid dendritic cells are not required to clear bone metastases in Ron SF$^{-/-}$ hosts. A**, Representative flow plot (left) and quantification (right) of DX5+NKp46+ peripheral blood NK cell frequency in Ron SF$^{-/-}$ mice following anti-NK1.1 depletion ($n$ = 4-8/group). **B**, Representative flow plot (left) and quantification (right) of Siglech+ bone marrow pDC frequency in Ron SF$^{-/-}$ mice following anti-BST2 depletion ($n$ = 7-10/group). **C**, Representative EpCAM IHC images of bones from NK1.1+ cell-depleted WT and Ron SF$^{-/-}$ mice, 4 weeks post-IT injection (left), with quantification of the number of mice with tumors (right; $n$ = 9 WT, 15 Ron SF$^{-/-}$). Scale bars, 1 mm. **D**, Representative EpCAM IHC images of bones from BST2+ cell-depleted WT and Ron SF$^{-/-}$ mice, 4 weeks post-IT injections (left), with numerical analysis of the presence of tumors (right; $n$ = 16 WT, 18 Ron SF$^{-/-}$). Scale bars, 1 mm. $P$ values were calculated using one-way ANOVA (**A** and **B**) and Fisher's exact test (**C** and **D**). ns = not significant, $P \geq 0.05$; *, $P < 0.05$; **, $P < 0.01$; ***, $P < 0.001$; ****, $P < 0.0001$.

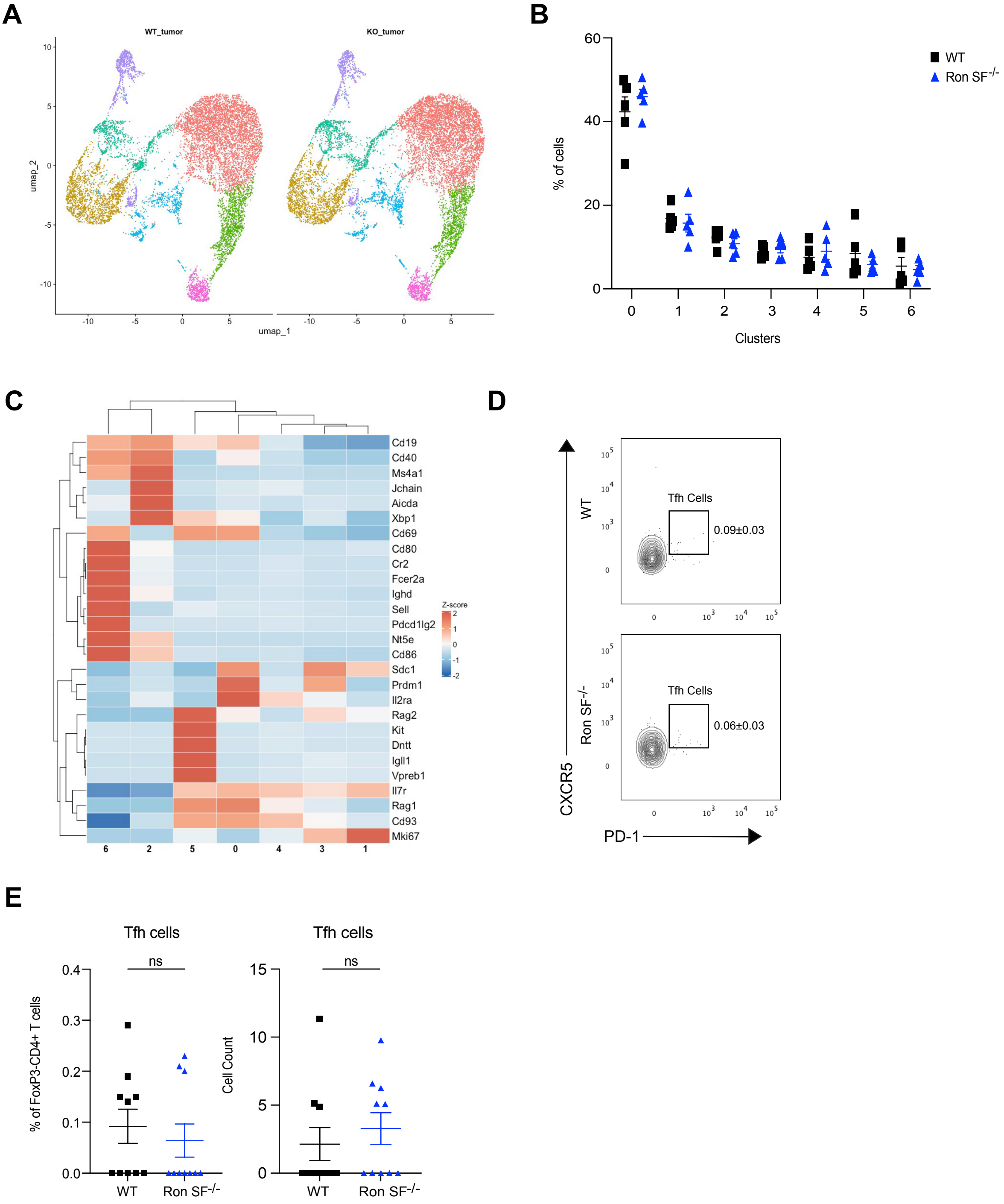
A
WT_tumor
KO_tumor
umap_2
umap_1
B
% of cells
WT
Ron SF-/-
Clusters
C
Z-score
Cd19
Cd40
Ms4a1
Jchain
Aicda
Xbp1
Cd69
Cd80
Cr2
Fcer2a
Ighd
Sell
Pdcd1lg2
Nt5e
Cd86
Sdc1
Prdm1
Il2ra
Rag2
Kit
Dntt
Igll1
Vpreb1
Il7r
Rag1
Cd93
Mki67
D
WT
Tfh Cells
0.09±0.03
Ron SF-/-
Tfh Cells
0.06±0.03
CXCR5
PD-1
E
Tfh cells
ns
% of FoxP3-CD4+ T cells
WT
Ron SF-/-
Tfh cells
ns
Cell Count
WT
Ron SF-/-

**Fig. S4**. **scRNA sequencing of B220+ cells from tumor-enriched areas of WT and Ron SF$^{-/-}$ bones.** **A**, Individual UMAPs from each group. **B**, Proportions of cells in each cluster in each genotype (*n* = 5/group). **C**, Heatmap showing the expression of representative genes from different B cell subsets, comparing each cluster against the remaining clusters. The color scale represents normalized expression by Z scores. **D**, Representative flow plots of CXCR5+PD1+ bone marrow Tfh cell frequency in WT and Ron SF$^{-/-}$ tumor naïve mice. **E**, Quantification of CXCR5+PD1+ bone marrow Tfh cell frequency and cell count in WT and Ron SF$^{-/-}$ tumor naïve mice (n = 10/group). Data are represented as mean ± SEM. *P* values were calculated using an unpaired two-tailed Student *t* test. ns = not significant.

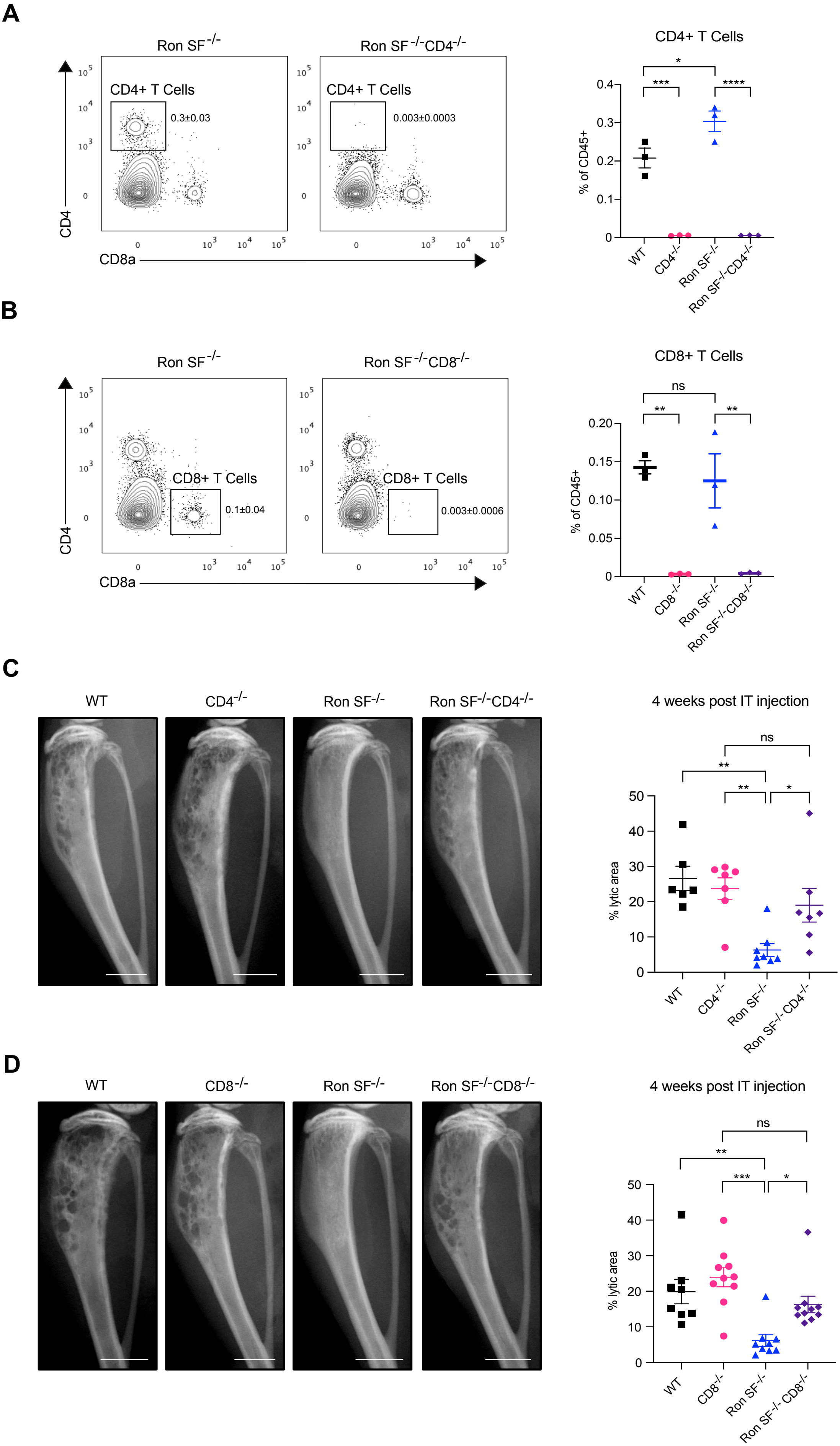
A
Ron SF-/-
Ron SF-/-CD4-/-
CD4+ T Cells
0.3±0.03
0.003±0.0003
CD4
CD8a
CD4+ T Cells
% of CD45+
WT
CD4-/-
Ron SF-/-
Ron SF-/-CD4-/-
B
Ron SF-/-
Ron SF-/-CD8-/-
CD8+ T Cells
0.1±0.04
0.003±0.0006
CD4
CD8a
CD8+ T Cells
% of CD45+
WT
CD8-/-
Ron SF-/-
Ron SF-/-CD8-/-
ns
C
WT
CD4-/-
Ron SF-/-
Ron SF-/-CD4-/-
4 weeks post IT injection
% lytic area
ns
D
WT
CD8-/-
Ron SF-/-
Ron SF-/-CD8-/-
4 weeks post IT injection
% lytic area
ns

**Fig. S5. Loss of CD4+ T cells and CD8+ T cells results in osteolytic lesions in Ron SF$^{-/-}$ hosts.** **A**, Representative flow plot (left) and quantification (right) of CD3e+CD4+ bone marrow T cell frequency in Ron SF$^{-/-}$ and Ron SF$^{-/-}$CD4$^{-/-}$ mice (*n* = 3/group). **B**, Representative flow plot (left) and quantification (right) of CD3e+CD8+ bone marrow T cell frequency in Ron SF$^{-/-}$ and Ron SF$^{-/-}$CD8$^{-/-}$ mice (*n* = 3/group). **C**, Representative X-rays of WT, CD4$^{-/-}$, Ron SF$^{-/-}$, and Ron SF$^{-/-}$CD4$^{-/-}$ bones (left) and lytic area quantification (right), 4-weeks post-IT injection (*n* = 6-7/group). Scale bars, 2 mm **D**, Representative X-rays of WT, CD8$^{-/-}$, Ron SF$^{-/-}$, and Ron SF$^{-/-}$CD8$^{-/-}$ bones (left) and lytic area quantification (right) at the 4-week timepoint (*n* = 8-10/group). Scale bars, 1 mm. *P* values were calculated using one-way ANOVA. ns = not significant, $P \geq 0.05$; *, $P < 0.05$; **, $P < 0.01$; ***, $P < 0.001$; ****, $P < 0.0001$.

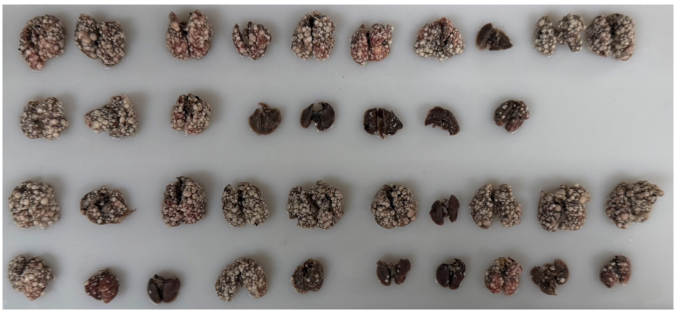

WT

Ron SF$^{-/-}$

JH$^{-/-}$

Ron SF$^{-/-}$JH$^{-/-}$

**Fig. S6**. **Loss of mature B cells does not rescue tumor growth in the lungs of Ron SF$^{-/-}$ hosts.** Image of gross metastatic outgrowth in WT, JH$^{-/-}$, Ron SF$^{-/-}$, and Ron SF$^{-/-}$JH$^{-/-}$ lungs.
